# Antibody Conversion rates to SARS-CoV-2 in Saliva from Children Attending Summer Schools in Barcelona, Spain

**DOI:** 10.1101/2021.04.20.440593

**Authors:** Carlota Dobaño, Selena Alonso, Mariona Fernández de Sevilla, Marta Vidal, Alfons Jiménez, Gemma Pons Tomas, Chenjerai Jairoce, María Melé Casas, Rocío Rubio, María Hernández García, Gemma Ruiz-Olalla, Mònica Girona-Alarcón, Diana Barrios, Rebeca Santano, Robert A. Mitchell, Laura Puyol, Leonie Mayer, Jordi Chi, Natalia Rodrigo Melero, Carlo Carolis, Aleix Garcia-Miquel, Elisenda Bonet-Carne, Joana Claverol, Marta Cubells, Claudia Fortuny, Victoria Fumadó, Cristina Jou, Carmen Muñoz-Almagro, Luis Izquierdo, Quique Bassat, Eduard Gratacós, Ruth Aguilar, Juan José García-García, Gemma Moncunill, Iolanda Jordan

**Affiliations:** ISGlobal, Hospital Clínic - Universitat de Barcelona, Barcelona, Catalonia, Spain; Consorcio de Investigación Biomédica en Red de Epidemiología y Salud Pública (CIBERESP), Madrid, Spain; Institut de Recerca Sant Joan de Déu, Esplugues, Barcelona, Spain; Pediatrics Department, Hospital Sant Joan de Déu, Universitat de Barcelona, Esplugues, Barcelona, Spain; Paediatric Intensive Care Unit, Hospital Sant Joan de Déu, Universitat de Barcelona, Barcelona, Spain; Biomolecular screening and Protein Technologies Unit, Centre for Genomic Regulation (CRG), The Barcelona Institute of Science and Technology, Barcelona, Spain; Fetal Medicine Research Center (Hospital Clínic and Hospital Sant Joan de Déu), Universitat de Barcelona, Barcelona, Spain; Institut d’Investigacions Biomèdiques August Pi i Sunyer (IDIBAPS), Barcelona, Spain; Universitat Politècnica de Catalunya, BarcelonaTech, Spain; Fundació Sant Joan de Déu, Barcelona, Spain; Infectious Diseases Department, Hospital Sant Joan de Déu, Barcelona, Spain; Department of Pathology and Biobank Hospital Sant Joan de Déu, Barcelona, Spain; CIBERER, Instituto de Salud Carlos III, Barcelona, Spain; Department of Medicine, Universitat Internacional de Catalunya, Barcelona, Spain; Molecular Microbiology Department, Hospital Sant Joan de Déu, Esplugues, Barcelona, Spain; Centro de Investigação em Saúde de Manhiça (CISM), Maputo, Mozambique; ICREA, Pg. Lluís Companys 23, 08010 Barcelona, Spain; Center for Biomedical Research on Rare Diseases (CIBER-ER), Madrid, Spain

**Keywords:** SARS-CoV-2, antibody conversion, saliva, children, schools

## Abstract

Surveillance tools to estimate infection rates in young populations are essential to guide recommendations for school reopening and management during viral epidemics. Ideally, field-deployable non-invasive, sensitive techniques are required to detect low viral load exposures among asymptomatic children. We determined SARS-CoV-2 antibody conversion by high-throughput Luminex assays in saliva samples collected weekly in 1,509 children and 396 adults in 22 Summer schools and 2 pre-schools in 27 venues in Barcelona, Spain, from June 29^th^ to July 31^st^ 2020, between the first and second COVID-19 pandemic waves. Saliva antibody conversion defined as ≥4-fold increase in IgM, IgA and/or IgG levels to SARS-CoV-2 antigens between two visits over a 5-week period was 3.22% (49/1518), or 2.36% if accounting for potentially cross-reactive antibodies, six times higher than the cumulative infection rate (0.53%) by weekly saliva RT-PCR screening. IgG conversion was higher in adults (2.94%, 11/374) than children (1.31%, 15/1144) (p=0.035), IgG and IgA levels moderately increased with age, and antibodies were higher in females. Most antibody converters increased both IgG and IgA antibodies but some augmented either IgG or IgA, with a faster decay over time for IgA than IgG. Nucleocapsid rather than spike was the main antigen target. Anti-spike antibodies were significantly higher in individuals not reporting symptoms than symptomatic individuals, suggesting a protective role against COVID-19. To conclude, saliva antibody profiling including three isotypes and multiplexing antigens is a useful and more user-friendly tool for screening pediatric populations to determine SARS-CoV-2 exposure and guide public health policies during pandemics.

## INTRODUCTION

Children infected with the severe acute respiratory syndrome coronavirus (SARS-CoV-2) usually present milder forms of the coronavirus disease (COVID-19) or are often asymptomatic, although they seem to be similarly susceptible to getting infected and can therefore transmit the virus to other people (1–4). Paradoxically, they are regarded as highly vulnerable to the “collateral damage” of COVID-19, i.e. huge social, family, economic and mental health foreseen consequences (5). The lack of attention to this specific age group has generated stress and has prevented evidence-based information to guide public health policies specifically designed for this population.

From a social perspective, there is an urgent need to have solid data on how COVID-19 affects children, and what is the contribution of this age group to overall community transmission. This implies the need to better define the patterns of SARS-CoV-2 transmission in pediatric populations, how many acquire the virus and whether they can infect other children and/or adults (4,6–8). Research oriented to answer these questions offers unique opportunities to raise evidence to support policies regarding maintaining schools and extra-curricular activities open.

Several studies have investigated the epidemiology and clinical presentation of SARS-CoV-2 infection during the pandemic (1,9). Mostly, they report that children are probably diagnosed less often with COVID-19, but still there exist confounding factors and controversial reports (10,11). Several hypotheses have been postulated to explain the milder presentation of COVID-19 in children, including a putative protective role of pre-existing cross-reactive antibodies to common cold human coronaviruses (HCoV) (12,13), lower expression of ACE2 (14), and lower pro-inflammatory propensity in their immune system (15).

The different diagnostic procedures implemented to date for SARS-CoV-2 infection in children are essentially the same as those in adults. Nasopharyngeal swabs for real time polymerase chain reaction (RT-PCR) or protein antigen diagnosis are the preferred because of their higher sensitivity and specificity (16). To reduce the inconvenience and discomfort associated with nasopharyngeal samples, nasal swabs have also been approved (17). In addition, non-invasive and better accepted saliva sampling for RT-PCR has shown similar results to nasopharyngeal swabs (18).

However, such methods are only useful for diagnosis of a current infection, but are not able to establish the percentage of the population that has been exposed to SARS-CoV-2. For this purpose, antibody-based detection methods are more appropriate, given that certain antibody responses persist over time. Furthermore, antibody surveillance could increase the sensitivity to detect incidence of new cases in longitudinal cohorts by assessing antibody conversion rates in prospective samples, particularly among asymptomatic children who may have lower viral loads and possibly more frequent false negatives for RT-PCR and/or for antigen detection tests.

Antibody assays are usually performed using plasma or serum samples and can be done in saliva samples (19–21), although they are not implemented in clinical practice. They offer many logistic advantages over tests requiring blood samples, especially in pediatric patients and large studies. Versatile multiplex antibody assays measuring several isotypes (IgM, IgA, IgG) and multiple SARS-CoV-2 antigens (22), rather than only 1-2 antibodies/antigens in the commercial kits, offer the greatest sensitivity to detect and accurately quantify a breadth of specificities, increasing the potential to identify recently and past exposed individuals, even if they have lower antibody levels, e.g. in asymptomatic subjects. In addition, IgA plays a very important role in COVID-19 immunity (23) and interrogating saliva samples can shed more light into mechanisms of mucosal protection.

The primary objective of this study was to determine SARS-CoV-2 exposure and antibody conversion in two consecutive saliva samples, as a proxy of seroconversion and seroprevalence, in children and adult populations in a school-like environment, between the first and second COVID-19 pandemic waves in Barcelona, Spain.

## RESULTS

The characteristics of the participants tested for saliva antibodies are summarized in **Table 1**. Detailed baseline characteristics of the cohort and the incidence of RT-PCR infections are reported elsewhere (24). During the 5-week study period, the laboratory received and processed 5,368 saliva samples collected in 1,509 children and 396 adult participants. In this cohort, only 5 adults had a prior diagnosis of COVID-19. For the antibody determinations, 3,475 inactivated saliva samples with sufficient volume, including first and last paired visits and those with only one visit available, were analyzed. Mean time between first and last visits was 15.69 days (SD 6.44). Between those two visits, 7 children and 3 adults tested positive for SARS-CoV-2 RT-PCR.

**Table 1.**
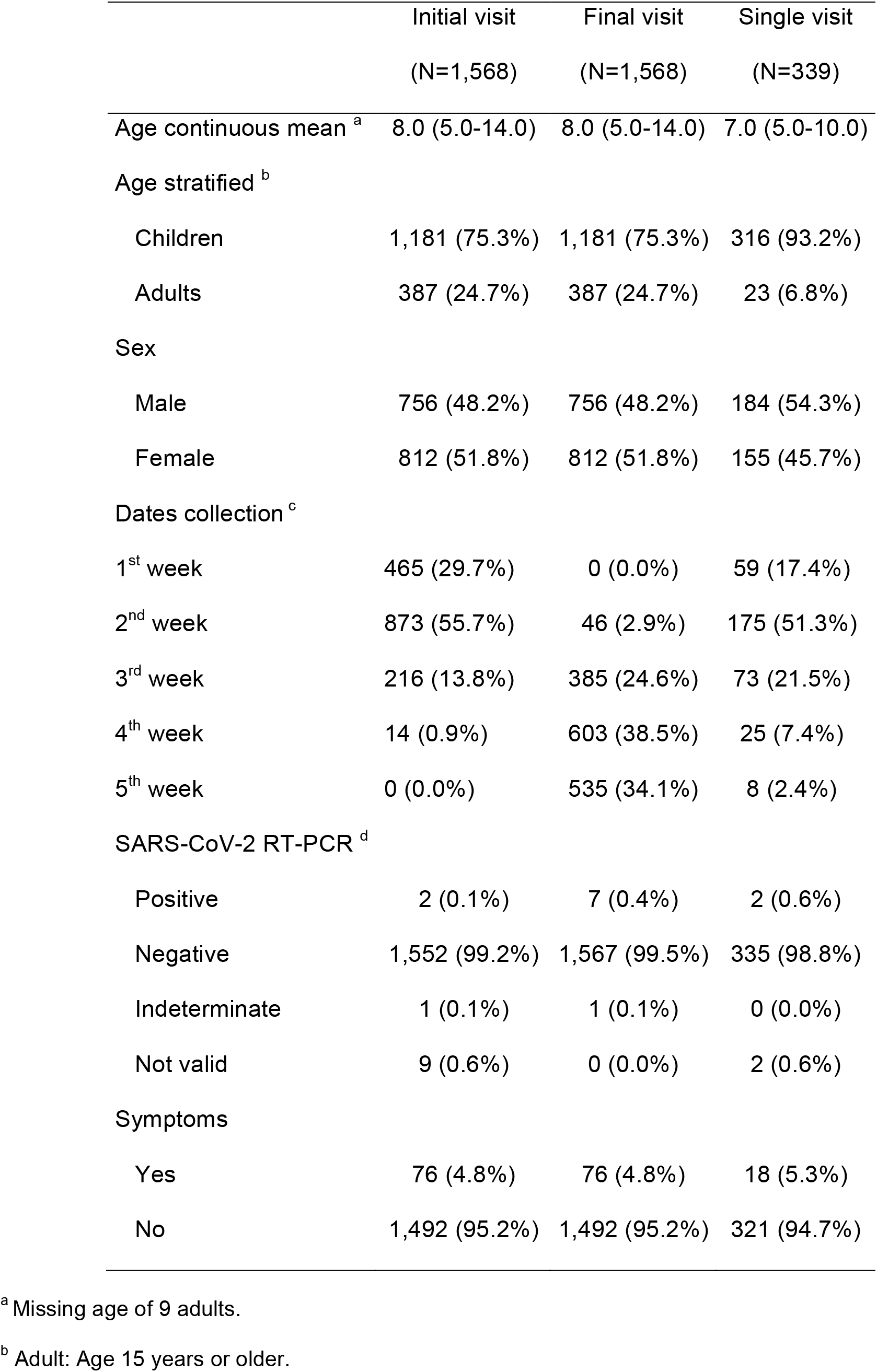
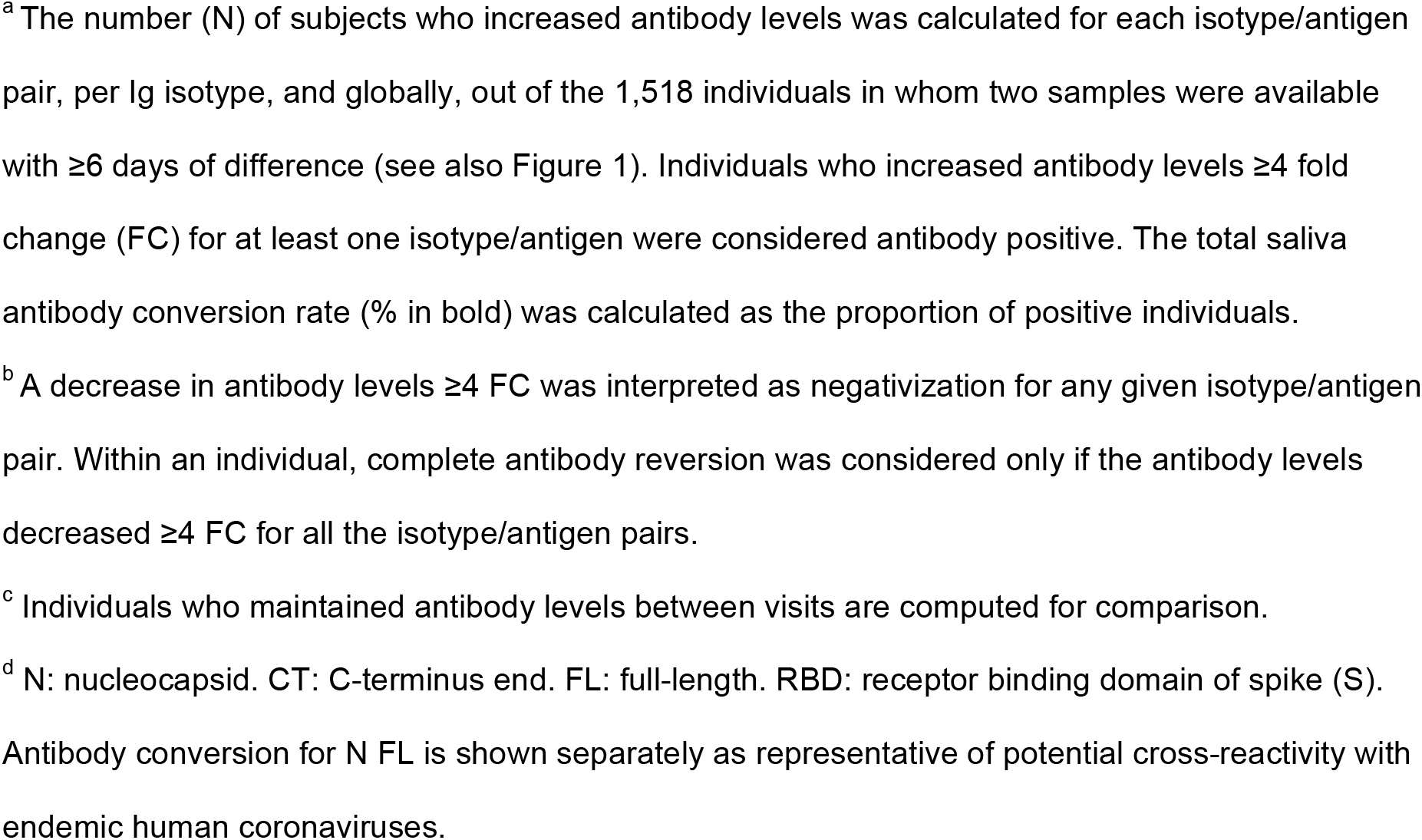
Characteristics of participants in the Summer school longitudinal study. All individuals with at least one saliva sample of sufficient volume available are included.

### SARS-CoV-2 antibody saliva conversion

To quantify the number of subjects acquiring an infection, we first identified those in whom saliva antibody levels to SARS-CoV-2 changed ≥3-4-fold from the first to the last visit (**Figures 1 & S1**). Computing the individuals with a ≥4-fold increase in antibody levels to at least one Ig/antigen pair, the overall antibody conversion rate was 3.22% (49/1518) (**Table 2**). This represented a 6 times higher estimate of new SARS-CoV-2 infections than what RT-PCR detected in this subgroup (8/1518, 0.53%). **Figure S2** shows the antibody levels in those diagnosed as RT-PCR positive in the study cohort. Stratifying by age, antibody conversion rates for IgG between the first and last visit were significantly higher in adults (2.94%, 11/374) than in children (1.31%, 15/1144) (p=0.035) (**Table S1**). Antibody conversion was higher for IgA (2.37%, 36/1518) and IgG (1.71%, 26/1518) than for IgM (0.2%, 3/1518). The N FL and N CT proteins were the main targets of saliva antibodies, followed by the S2 protein. Excluding individuals who only increased SARS-CoV-2 N FL antibodies (0.86%, 13/1518), potentially cross-reactive with HCoV N FL (25), the adjusted conversion rate was 2.36%. Antibodies were maintained at a wide range of levels in a large number of subjects, and in others they decayed from the first to the last visit (**Table 2, Figure 1**), but no one reverted for all isotypes/antigens (≥4-fold decrease). IgA to SARS-CoV-2 antigens reverted more than IgG antibodies. Some subjects maintaining antibodies (**Figure 1**), or with only one sample collection (**Figure S3**), had high levels similar to those classified as positive among the subjects who converted (last visit) or reverted (first visit). We explored the prevalence of SARS-CoV-2 exposure in all individuals using FMM and EM algorithms (**Figure S4**). Although the models correctly classified the saliva antibody converters as positive, the population positivity estimates obtained at each time point were substantially higher than expected (**Table S2**).

**Table 2.**
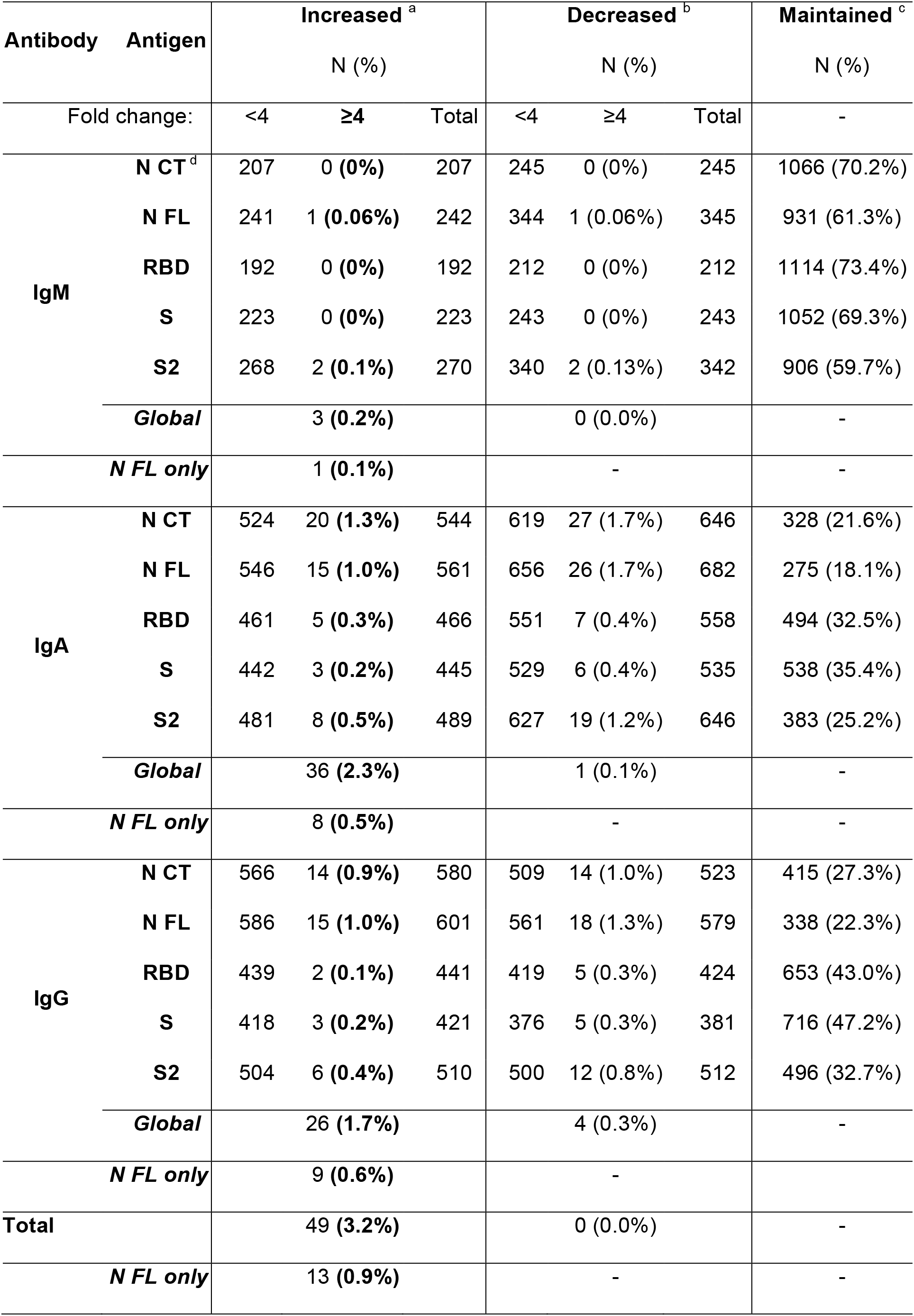
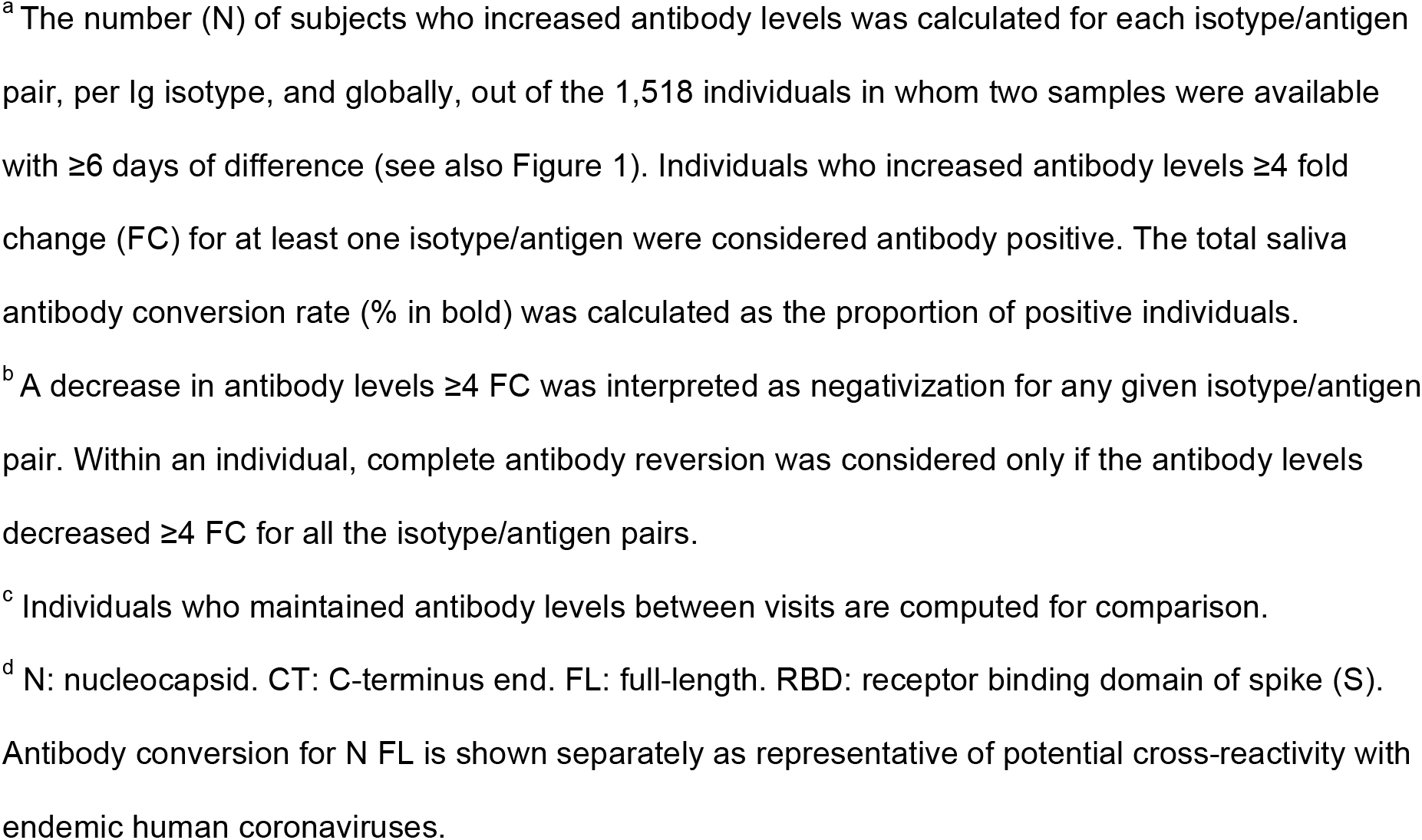
Saliva antibody conversion rates between the first and last study visit.

**Figure 1.**
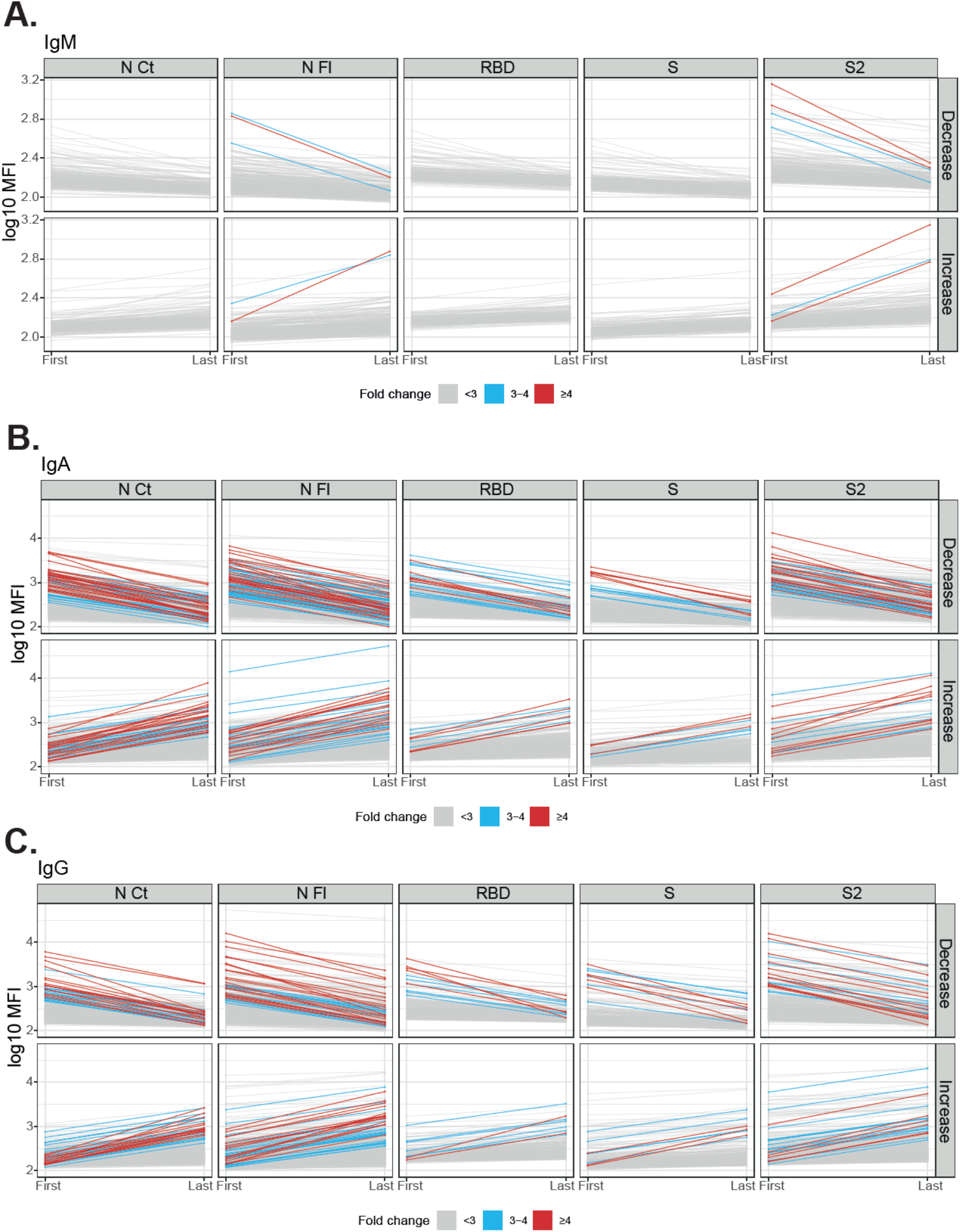
Evolution of IgM, IgA and IgG levels to SARS-CoV-2 antigens between the first and last visit in paired samples. Individuals who decreased or increased IgM (A), IgA (B) or IgG (C) levels per each isotype and antigen are shown in different plots. Grey lines mean <3 fold-change, blue lines mean 3-4-fold change, and red lines mean ≥4 fold-change. Table 1 indicates the number and proportion of individuals in each category. The levels of antibodies in individuals with only one sample are depicted in Figure S3.

### Factors affecting SARS-CoV-2 antibody levels

Levels of IgA and IgG antibodies in saliva were significantly lower in children (n=2,677) than in adults (n=800), while no differences were seen for IgM (**Figure 2A**). IgG and IgA levels gradually increased statistically significantly with age (**Figure S5**). Among RT-PCR positives (n=10), IgA and IgG levels tended to also be higher in adults compared to children (**Figure S6**). Patients with COVID-19 compatible symptoms had statistically significantly lower antibody levels to S, S2 and RBD than individuals not reporting symptoms (**Figure 2B**). Levels of antibodies of the three isotypes were higher in females than males (**Figure S7**).

**Figure 2.**
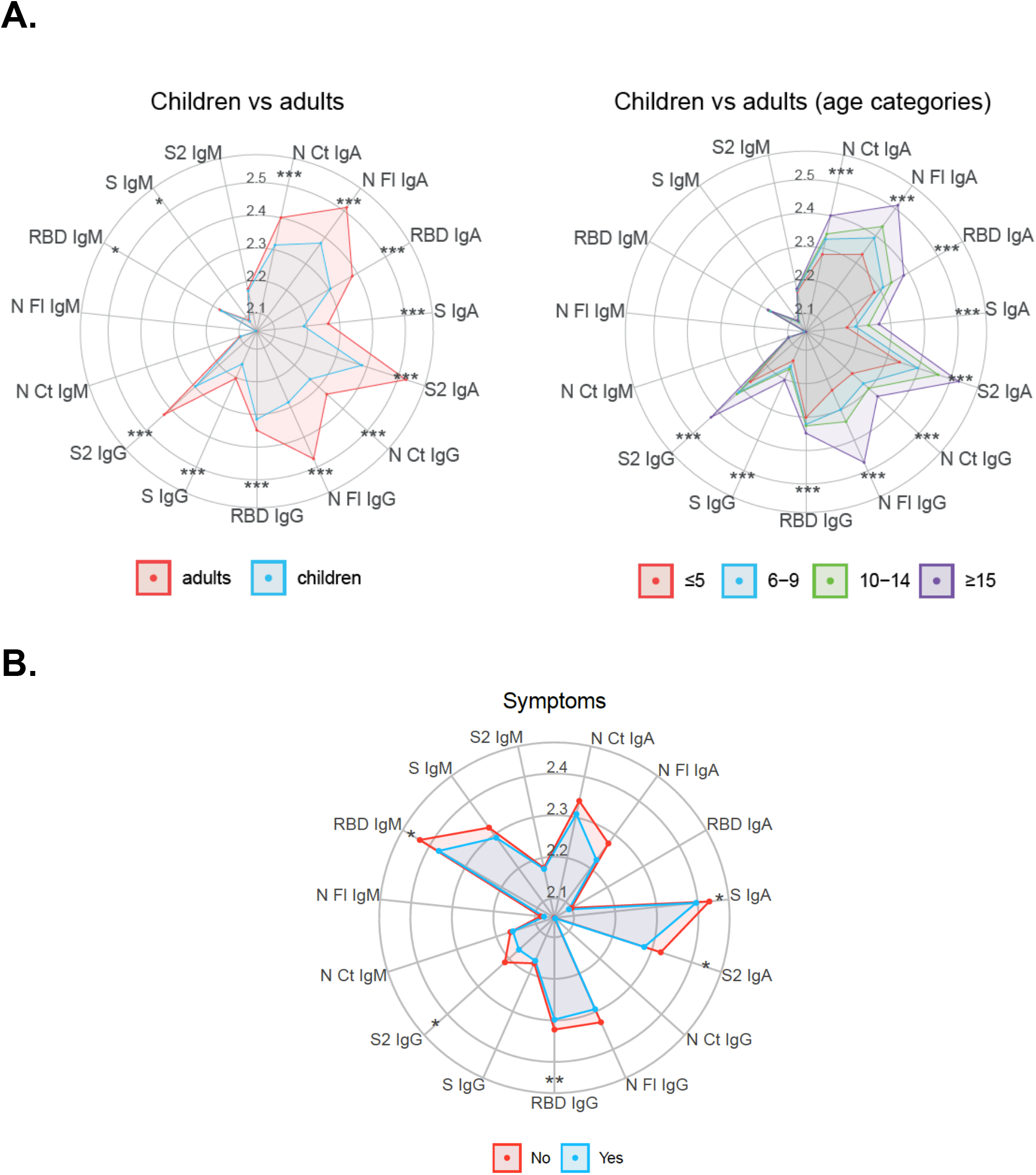
Antibody levels by age and symptoms. Radar charts of median antibody levels in the last and single visits comparing children (<15 years old) versus adults **(A)**. Median antibody levels comparing symptomatic (n=52, blue) versus asymptomatic (n=3,423, red) individuals (**B**). Group medians were compared through Mann-Whitney U test. * p ≤ 0.05, ** p ≤ 0.01, *** p ≤ 0.001.

### Multi-marker antibody response patterns

In a heatmap of FC antibody responses from first to last visit, most individuals with high FC showed an increase in both IgG and IgA antibodies (very few IgM), and a smaller group showed an increase in either IgG or IgA antibodies only (**Figure 3A**). Focusing on individuals with ≥4-FC in levels between visits, some increased IgG predominantly, some increased IgA predominantly, and others increased both isotypes (**Figure 3B**). There was no clear pattern for age or symptoms.

**Figure 3.**
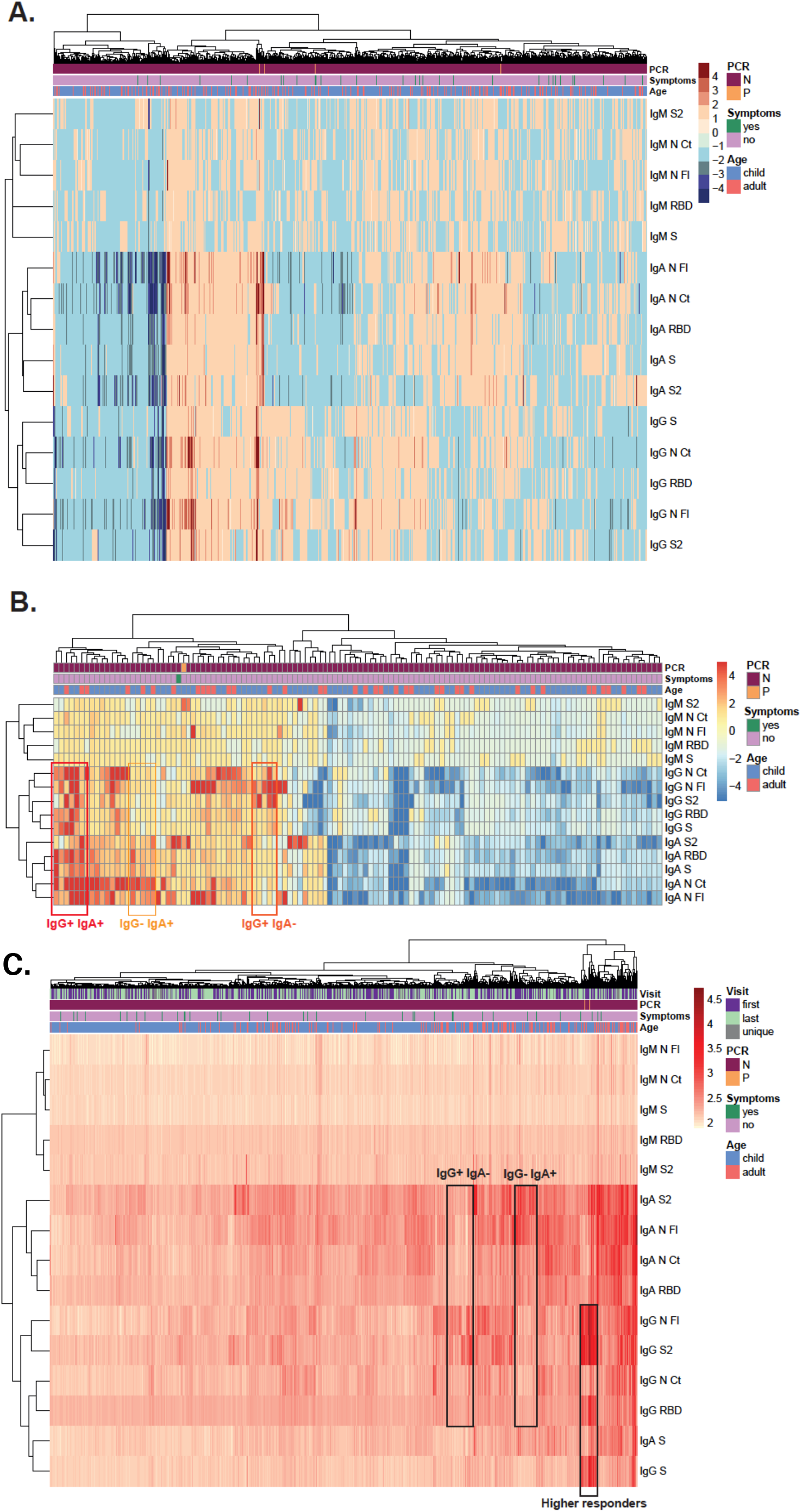
Heatmap analysis of antibody responses per individual. Fold change antibody levels (MFI) with hierarchical clustering (Canberra), including all individuals with paired first and last visit samples, showing decreasers (blue scale), maintainers and increasers (red scale) **(A)** or including only individuals who increased or decreased antibody levels ≥4-fold between the two visits **(B)**. Antibody levels (log_10_ MFI) with hierarchical clustering (Euclidean) in all individuals (**C**).

Combining all antibodies and variables in all individuals, the strongest signal for the high responders mapped to IgG to N FL and S2 antigens, as seen in the antibody conversion analysis, which was accompanied by IgG to S and RBD responses, but lower IgA reactivity (**Figure 3C**). Another group of antibody responders had a more predominant IgA than IgG reactivity, while others had a more predominant IgG than IgA reactivity. There were more adults among higher antibody responders and no clear pattern was seen for symptoms.

## DISCUSSION

We showed that a non-invasive screening approach based on weekly saliva sampling in ~2000 subjects with thousands of visits, coupled to a high-throughput multiplex assay to quantify antibodies, is capable of measuring infection rates in pediatric populations with high sensitivity. Thus, saliva antibody conversion between two study visits over a 5-week period in our population, based on a ≥4-FC increase combining 3 immunoglobulin isotypes and 5 SARS-CoV-2 antigens, was 3.22%, or 2.36% excluding individuals with only N FL antibodies that may cross-react between SARS-CoV-2 and HCoV (25). Interestingly, saliva IgG conversion rates, and levels of IgA and IgG, were significantly lower in children than adults, consistent with a differential infection and transmission dynamics. In addition to circumventing the need for blood sampling, saliva surveys are easier to deploy in the field and do not require qualified health care personnel for collection.

Saliva antibody conversion estimates were 6 times higher than the cumulative infection rate derived from weekly RT-PCR screening, despite capturing exposure to the virus with ~10-14 days delay in respect to the infection, and that peak IgG levels are usually attained ~16-30 days post symptoms onset. Of note, 6 out of 8 RT-PCR positive individuals had the viral diagnosis at the final visit, therefore we would not expect antibody conversion in those until some days later. The finding that a number of potential infections were detected by saliva antibody FC but not by RT-PCR, could be related to lower viral loads in asymptomatics and in children (the predominant population here), consistent with their lower antibody levels compared to adults. Another explanation for possible RT-PCR false negatives is that the virus presence could be more transitory in children. The discrepancy would not be explained by the presence of high antibody levels at the first or only visit, suggesting previous exposure before the Summer school, because only those with low antibody levels at the initial visit would contribute to the ≥4-FC conversion. Importantly, other studies have also shown that children with a negative RT-PCR can have antibody responses detectable in saliva (26). Together, data indicate that children can mount an antibody response to SARS-CoV-2 without viral diagnosis, suggesting that immunity in children could prevent the establishment of SARS-CoV-2 infection.

Further supporting a role for saliva antibodies on immunity, mucosal IgM, IgA and IgG to S but not N proteins were significantly higher in individuals not reporting symptoms than in symptomatic ones. This is the opposite of what is commonly observed in blood: symptomatic or severe disease patients have higher viral loads and SARS-CoV-2 antibody levels than asymptomatic individuals, reflecting intensity of exposure. Our results point to an anti-disease effect of saliva S-specific antibodies that are known to neutralize SARS-CoV-2 invasion via ACE2 receptors in respiratory mucosal tissues, being S the component of efficacious COVID-19 vaccines. Indeed, there is increasing data on the significant role for mucosal immunity and particularly for secretory as well as circulating IgA antibodies in COVID-19 (27). Mucosal IgA can have a key role in early SARS-CoV-2-specific neutralizing response (23). Patients with high saliva viral loads developed antiviral antibodies later than those with lower viral loads (27). Therefore, studies detecting IgA in addition to IgG in saliva will help to better understand the dynamics of COVID-19 mucosal immunity.

Due to the more transient nature of SARS-CoV-2 specific antibody responses in oligosymptomatic patients, reliance on measuring serum IgA and IgG might underestimate the percentage of individuals who have experienced COVID-19. In addition to serum, measurement of mucosal IgA should be considered, as local responses may be higher than systemic in such cases, or it could be that the response is only mucosal. IgA in mild COVID-19 cases can often be transiently positive in serum (28), and serum IgG may remain negative or become positive many days after symptom onset, while IgA could appear faster in saliva. Thus, an added benefit of saliva serological surveys is that, in people with no or transient IgA or IgG serum responses but detectable IgA levels in nasal fluid and tears, exposure might only be easily detected in saliva but not blood antibodies (28). In our study, the measurement of both IgG and IgA in saliva increased the probability to identify positive responders because not all subjects produced both isotypes at the time of sampling.

Regarding antibody kinetics, many individuals appeared to maintain Ig levels similar to the ones observed in increasers over the follow up period, with no reversions. A faster decay in antibodies was seen for IgA than for IgG, consistent with its shorter half-life. Most studies show that systemic IgG antibodies are maintained in a majority of COVID-19 patients for at least 8-9 months post symptoms onset (29–31). Less information is available on the long-term kinetics of mucosal antibodies. Acknowledging that antibody concentrations in saliva are much lower than those in plasma, it would be relevant to investigate the long-term kinetics in future follow up studies.

Levels of saliva antibodies were higher to N than to S antigens. This shows that antibodies to N proteins, which are not included in current first-generation vaccines, are nevertheless immunogenic and may be useful to track viral exposure in saliva field surveys and after vaccination. Our prior studies in plasmas found evidence of higher cross-reactivity for N than S antigens among different coronaviruses, and suggested that higher levels of pre-existing antibodies to some seasonal HCoV could provide partial immunity against COVID-19 (25,32).

The main study limitation was the unavailability of pre-pandemic saliva samples that did not allow establishing the positivity threshold with saliva negative controls by the classical methods. Thus, we explored less standard FMM and EM algorithms to estimate the overall prevalence of SARS-CoV-2 exposure in the population studied. Models estimated positivity ranges that were much higher than seroprevalences estimated in contemporaneous surveys in the same geographical area (33–35). These algorithms may greatly overestimate the Ig positivity in saliva samples and/or saliva samples may be more sensitive to detect exposure to SARS-CoV-2, and/or classical methods to calculate positivity in serum/plasma may underestimate seroprevalence. More work is required to distinguish between these potential explanations, and to refine and validate these statistical approaches with datasets that have appropriate negative and positive controls. A related constraint was that we could not relate the antibody responses in saliva to current infection because there were very few RT-PCR positives, and that we could not compare saliva to serum responses due to the unavailability of blood samples. Therefore, our data could not be contrasted with the seroconversion or seroprevalence (36), considered the ‘gold standard’.

In conclusion, antibody profiling in saliva samples with a multiplex technique represents a helpful and simpler tool in community-based surveys for determining saliva antibody conversion and prevalence of SARS-CoV-2 exposure in a school-like environment. Saliva antibodies and conversion were lower in children than adults, and levels were higher in asymptomatic than symptomatic individuals, pointing to an anti-disease protective role of mucosal immunoglobulins. This non-invasive screening technique can help studying the dynamics of the pandemic in children and guiding policies about maintaining schools and holiday camps active. This approach will also be useful to study reinfections over time as well as immunogenicity and persistence of immunity after COVID-19 vaccination at a larger scale, due to the distinct N and S antigen specificities evaluated.

## METHODS

### Design, subjects and samples

A cohort of 1,905 children (age 0-14 years old) attending 22 Summer schools and 2 pre-schools, and adult staff working at the same facilities, located in 27 different venues in the Barcelona metropolitan region, Spain, was followed up from June 29^th^ to July 31^st^ 2020. Symptomatic children were defined as those with acute respiratory infection including fever, cough, headache, gastrointestinal symptoms, rhinorrhea or nasal congestion, anosmia or ageusia, dyspnea and myalgia.

Saliva samples were collected weekly over a 5-week period with Oracol devices (Malvern, UK) for optimal harvesting of crevicular fluid enriched with serum antibodies (37,38), transported refrigerated to ISGlobal lab on the same day for centrifugation, heat inactivation (60°C, 30 min) and −20°C freezing, until preparation of 37 pre-assay 96-well plates (G080, Attendbio) for antibody analysis.

The study protocol was approved by Sant Joan de Déu Ethics Committee (PIC-140-20) and followed the recommendations of the Helsinki Declaration. All participants or their legal guardians provided written informed consent before study procedures started.

### Laboratory measurements

Saliva SARS-CoV-2 RT-PCR detection was performed as described (24,39,40). Levels of IgG, IgA and IgM against SARS-CoV-2 nucleocapsid (N) full-length (FL) and C-terminus (CT) (25), spike (S), S2, and RBD proteins, were measured by Luminex assays (22) (supplementary methods) in saliva samples diluted 1:10 in 384-well plates, with paired samples from the same individual run together. Pre-pandemic negative controls were not available. Samples were acquired on a Flexmap 3D xMAP® and median fluorescent intensities (MFI) were exported for each analyte using xPONENT.

### Data analysis

Non-parametric Mann-Whitney U tests were used in boxplots to compare levels (log_10_MFI) of each antibody/antigen pair between study groups. Radar plots were used to compare median MFIs of all antibody responses between study groups by Mann-Whitney U test, adjusting p-values for multiple comparison by Benjamini-Hochberg. Heatmaps with hierarchical clustering (by Euclidian or Canberra methods) were used to evaluate patterns of responses at the individual level. To evaluate how many participants got infected during the study period, we considered at least a 3-4-fold increase (FC) in antibody levels between two consecutive visits (20). To define the overall saliva antibody conversion rate (primary endpoint), we applied the more stringent threshold of ≥4-FC in antibody levels from the first to the last week visit only in the subset of individuals in whom at least two samples were collected ≥6 days apart. Saliva antibody reversion was defined as a ≥3-4 FC decrease in antibody levels from the first to the last week visit in the same subset of individuals. As an exploratory endpoint, we estimated the prevalence of SARS-CoV-2 exposure at each timepoint, including individuals who were visited only once and those who maintained antibody levels between visits. Two approaches were tested based on positivity cutoffs calculated by Finite Mixture Models (FMM) (41) or with models estimated by the Expectation-Maximization (EM) (29) algorithms (see Supplementary Methods). All data were managed and analysed using R software v4.0.3 (devtools (30), tidyverse (31), ggplot2 (42), pheatmap (43), mclust (44) and cutoff (41,45) packages).

## ACKNOWLEDGEMENTS

We thank the volunteers for their participation in the study, and Llorenç Quintó for statistical advice on the FMM. We are indebted to the “Biobanc de l’Hospital Infantil Sant Joan de Déu per a la Investigació” for sample and data procurement.

This work was supported by the Stavros Niarchos Foundation, Banco Santander and other private donors through the Kids Corona Platform (Institut de Recerca Sant Joan de Deu), and the Fundació Privada Daniel Bravo Andreu. G.M. was supported by the Departament de Salut, Generalitat de Catalunya (grant number SLT006/17/00109). L.I. work was supported by PID2019-110810RB-I00 grant from the Spanish Ministry of Science & Innovation. Development of SARS-CoV-2 reagents was partially supported by the National Institute of Allergy and Infectious Diseases Centers of Excellence for Influenza Research and Surveillance (contract number HHSN272201400008C). ISGlobal receives support from the Spanish Ministry of Science and Innovation through the “Centro de Excelencia Severo Ochoa 2019-2023” Program (CEX2018-000806-S), and support from the Generalitat de Catalunya through the CERCA Program.

## SUPPLEMENTARY MATERIAL

## SUPPLEMENTARY METHODS

### RT-PCR diagnosis

Saliva collected for SARS-CoV-2 RNA detection was introduced into micronic tubes for pathogen inactivation (Zymo DNA/RNA Shield Lysis Buffer™; Zymo Research, Freiburg, Germany). Tubes were processed using a TecanEvo 200 automated liquid handling system. RNA was extracted using the Quick-DNA/RNA Viral MagBead kit (Zymo Research) that was fully automated on a TecanDreamPrep NAP Workstation (Tecan Trading AG, Switzerland). The RT-PCR assays were conducted according to CDC-006-00019 CDC/DDID/NCIRD/ Division of Viral Diseases protocol released on 3/30/2020 and available at https://www.fda.gov/media/134922/download, which included the CDC-approved primers and probes for SARS-CoV-2 N1 or N2 genes and RNaseP human gene as internal control^1,2^. Primers and probes were purchased from IDT integrated technologies (qPCR probes - 2019-nCoV CDC EUA Kit). RT-PCR assays were performed at the Centre for Genomic Regulation (CRG) in Barcelona, Spain, and results were validated by a Clinical Microbiologist of Hospital Sant Joan de Déu.

### Measurement of antibodies

SARS-CoV-2 target antigens assays to measure IgG, IgA and IgM included the nucleocapsid (N) full-length (FL) and C-terminus (amino acid residues 340-416, CT), the spike (S) FL produced at CRG, S2 purchased from SinoBiological, and RBD donated by F. Krammer (Mount Sinai, NY). For the quantitative suspension array technology (qSAT) assay, protein-coupled magnetic microspheres (Luminex Corporation, Austin, TX) were added to a 384-well μClear^®^ flat bottom plate (Greiner Bio-One, Frickenhausen, Germany) in multiplex (2000 microspheres per analyte per well) in a volume of 90 μL of Luminex Buffer (1% BSA, 0.05% Tween 20, 0.05% sodium azide in PBS) using Integra Viaflo semi-automatic device (96/384, 384 channel pipette). Pools of plasmas from adults exposed to SARS-CoV-2 were used as positive controls in 2-fold, 8 serial dilutions starting at 1:12.5. Technical blanks consisting of Luminex Buffer and microspheres without samples were added in 4 wells to detect and adjust for non-specific microsphere signal. Saliva negative controls were not added due to the unavailability of pre-pandemic samples in spite of contacting national biobanks. Ten μL of each dilution of the positive control and test saliva samples were added to the 384-well plate using Assist Plus Integra device with 12 channels Voyager pipette (final saliva dilution of 1:10). Paired samples from the same individual were run on the same plate. Plates were incubated for 1 h at room temperature in agitation (Titramax 1000) at 900 rpm and protected from light. Then, the plates were washed three times with 200 μL/well of PBS-T (0.05% Tween 20 in PBS), using BioTek 405 TS (384-well format). Twenty five μL of goat anti-human IgG-phycoerythrin (PE) (GTIG-001, Moss Bio) diluted 1:400, goat anti-human IgA-PE (GTIA-001, Moss Bio) 1:200, or goat anti-human IgM-PE (GTIM-001, Moss Bio) 1:200 in Luminex buffer were added to each well and incubated for 30 min. Plates were washed and microspheres resuspended with 80 μL of Luminex Buffer, covered with an adhesive film and sonicated 20 seconds on a sonicator bath platform, before acquisition on a Flexmap 3D xMAP® instrument. At least 50 microspheres per analyte per well were acquired. Crude median fluorescent intensities (MFI) and background fluorescence from blank wells were exported for each analyte using the xPONENT software.

### Exploratory antibody positivity analysis

An antibody positivity threshold could not be calculated by the classical method (mean + 3 standard deviations [SD] of negative control samples) due to the unavailability of pre-pandemic saliva samples. Finite Mixture Models (FMM) and Expectation-Maximization (EM) algorithms allow the classification of samples into two populations: negative and positive, and both are based on the EM algorithm. Briefly, the EM algorithm is an unsupervised clustering algorithm that works in two steps. The first step is the Expectation step, where the data are assigned to the closest centroid of the two clusters. The second step is the Maximization step, where the centroids of the clusters are recalculated with the newly assigned data. These two steps are repeated until the model converges. The location parameter *mu* and the scale parameter *sigma* are estimated for both of these distributions (populations). Then, we used the parameters and information generated by FMM and EM models to estimate cutoffs by the classical method or by the probability method in the natural scale for MFI levels. The classical approach calculates the cutoff as the mean + 3 standard deviation of the estimated seronegative population, and the probability approach defines the cutoff based on the classification estimated for each sample. Antibody conversion was also estimated based on the proportion of individuals who were antibody-negative at their first and positive at their last visit, as defined by the FMM and EM algorithms.

All data were managed and analysed using R software v4.0.3 and its packages devtools and tidyverse. The ggplot2 and pheatmap packages were used to perform graphs. The mclust and cutoff packages were used for EM and FMM models.

## SUPPLEMENTARY TABLES

**Table S1.** Fold change antibody levels between first and last visit stratified by age. The total number of individuals with paired samples and ≥6 days between them was 1,518, with 1,144 children (age < 15 years) and 374 adults. Antibody conversion was calculated for increase fold change (FC) antibody levels ≥ 4 between visits per immunoglobulin isotype, considering any antigen, and globally, considering any isotype (IgM or IgA or IgG). Antibody reversion was calculated considering that all isotype/antigen pairs had to decrease FC antibody levels ≥ 4 between visits. P-values obtained with two-proportions z-test.

| Isotype | Antigen | Antibody conversion<br>(FC increase $\geq 4$) | | | Antibody reversion<br>(FC decrease $\geq 4$) | |
| --- | --- | --- | --- | --- | --- | --- |
|  |  | Children | Adults | p-value | Children | Adults |
| IgM | N CT | 0 (0.0%) | 0 (0.0%) | - | 0 (0.0%) | 0 (0.0%) |
|  | N FL | 1 (0.08%) | 0 (0.0%) | 0.567 | 1 (0.1%) | 0 (0.0%) |
|  | RBD | 0 (0.0%) | 0 (0.0%) | - | 0 (0.0%) | 0 (0.0%) |
|  | S | 0 (0.0%) | 0 (0.0%) | - | 0 (0.0%) | 0 (0.0%) |
|  | S2 | 2 (0.17%) | 0 (0.0%) | 0.418 | 2 (0.2%) | 0 (0.0%) |
|  | Global | 3 (0.26%) | 0 (0.0%) | 0.321 | 0 (0%) | 0 (0%) |
| IgA | N CT | 13 (1.1%) | 7 (1.9%) | 0.278 | 20 (1.8%) | 7 (1.9%) |
|  | N FL | 9 (0.8%) | 6 (1.6%) | 0.165 | 19 (1.7%) | 7 (1.9%) |
|  | RBD | 3 (0.26%) | 2 (0.5%) | 0.424 | 5 (0.4%) | 2 (0.5%) |
|  | S | 2 (0.17%) | 1 (0.3%) | 0.726 | 4 (0.3%) | 2 (0.5%) |
|  | S2 | 6 (0.5%) | 2 (0.5%) | 0.981 | 11 (1.0%) | 8 (2.1%) |
|  | Global | 24 (2.1%) | 11 (2.9%) | 0.345 | 1 (0.1%) | 0 (0.0%) |
| IgG | N CT | 8 (0.7%) | 6 (1.6%) | 0.112 | 6 (0.5%) | 8 (2.1%) |
|  | N FL | 9 (0.8%) | 6 (1.6%) | 0.165 | 11 (1.0%) | 7 (1.9%) |
|  | RBD | 2 (0.17%) | 0 (0.0%) | 0.418 | 1 (0.1%) | 4 (1.0%) |
|  | S | 2 (0.17%) | 1 (0.3%) | 0.728 | 2 (0.2%) | 3 (0.8%) |
|  | S2 | 4 (0.35%) | 2 (0.5%) | 0.620 | 7 (0.6%) | 5 (1.3%) |
|  | Global | 15 (1.3%) | 11 (2.9%) | 0.035 | 1 (0.1%) | 3 (0.8%) |
| **Total** |  | **34 (3.0%)** | **15 (4.0%)** | **0.323** | **0 (0%)** | **0 (0.0%)** |

N: nucleocapsid. FL: full-length. CT: C-terminus end. RBD: receptor binding domain of spike (S).

**Table S2.**
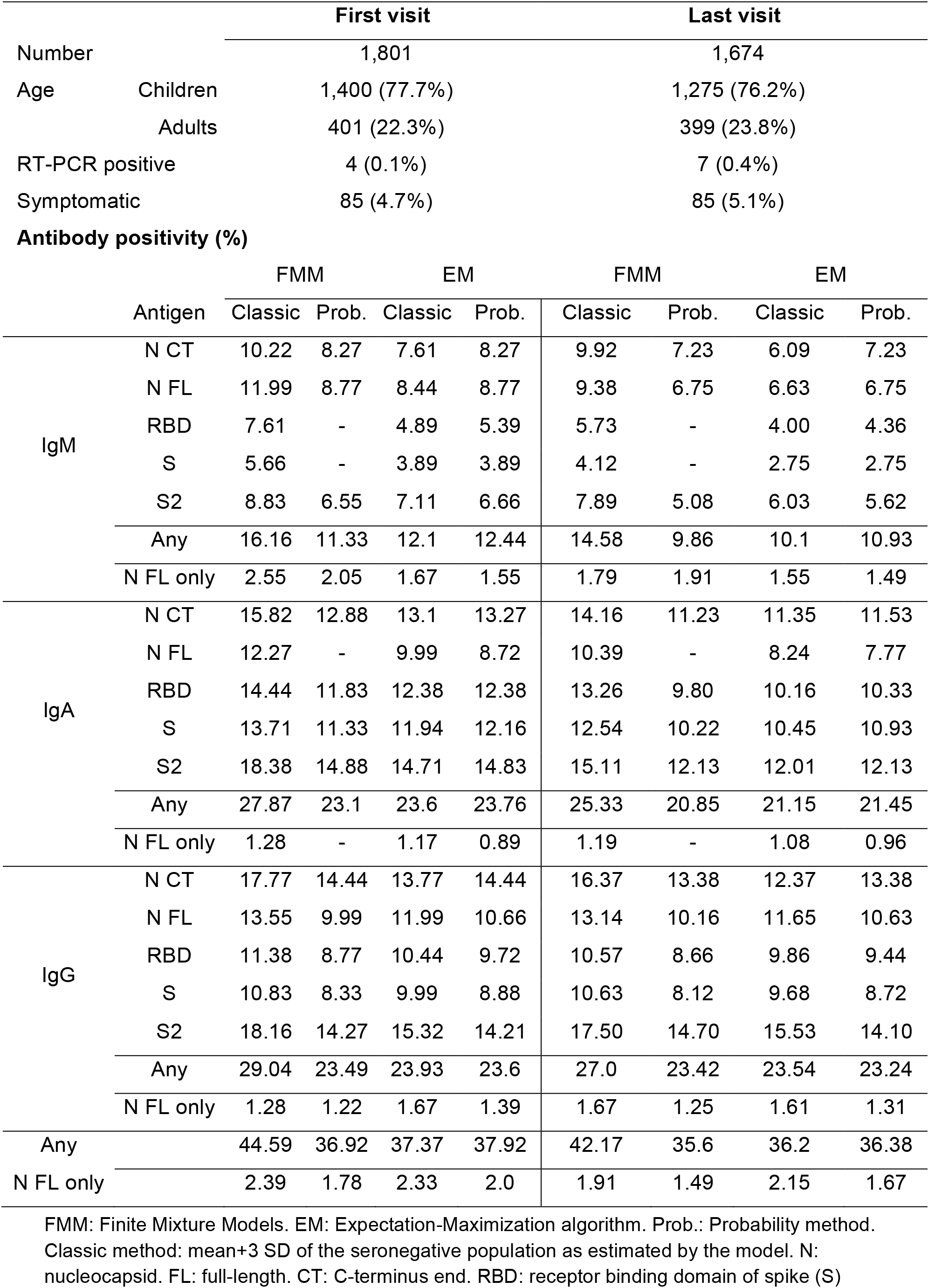
Percentage of positivity of each antibody and antigen per visit, including all saliva samples. See Figure S4 for graphical representation.

## SUPPLEMENTARY FIGURES

**Figure S1.**
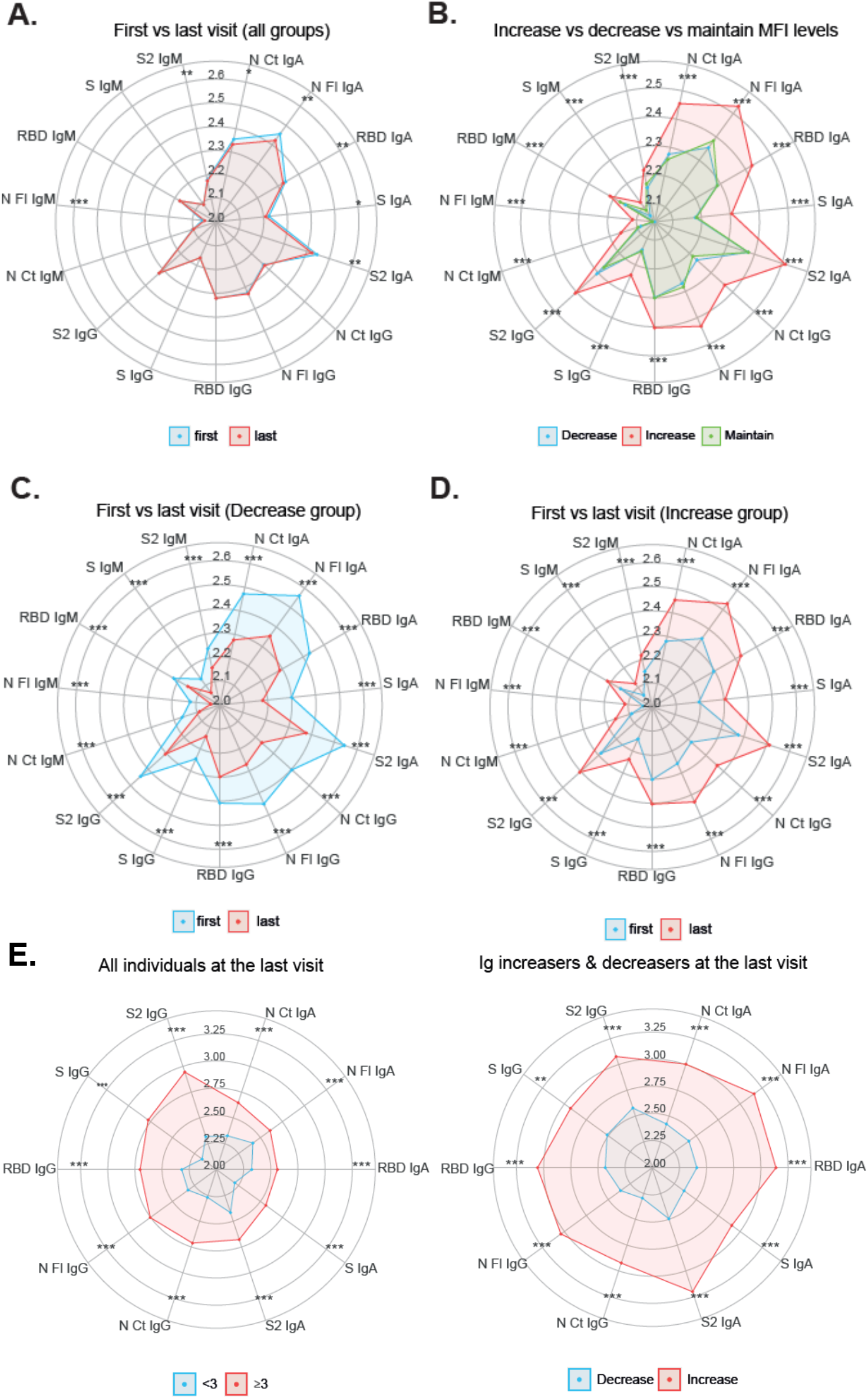
Radar charts of saliva antibodies by visit. Overall median antibody levels in the first and last visit **(A)**, in the last visit comparing individuals who increased, decreased (≥3 fold-change [FC]) or maintained (<3 FC) responses **(B)**, in the first versus the last visit in individuals who decreased **(C)** or increased **(D)** responses. FC in median antibody levels between first and last visit **(E)**. Groups were compared through Mann-Whitney test. * p ≤ 0.05, ** p ≤ 0.01, *** p ≤ 0.001.

**Figure S2.**
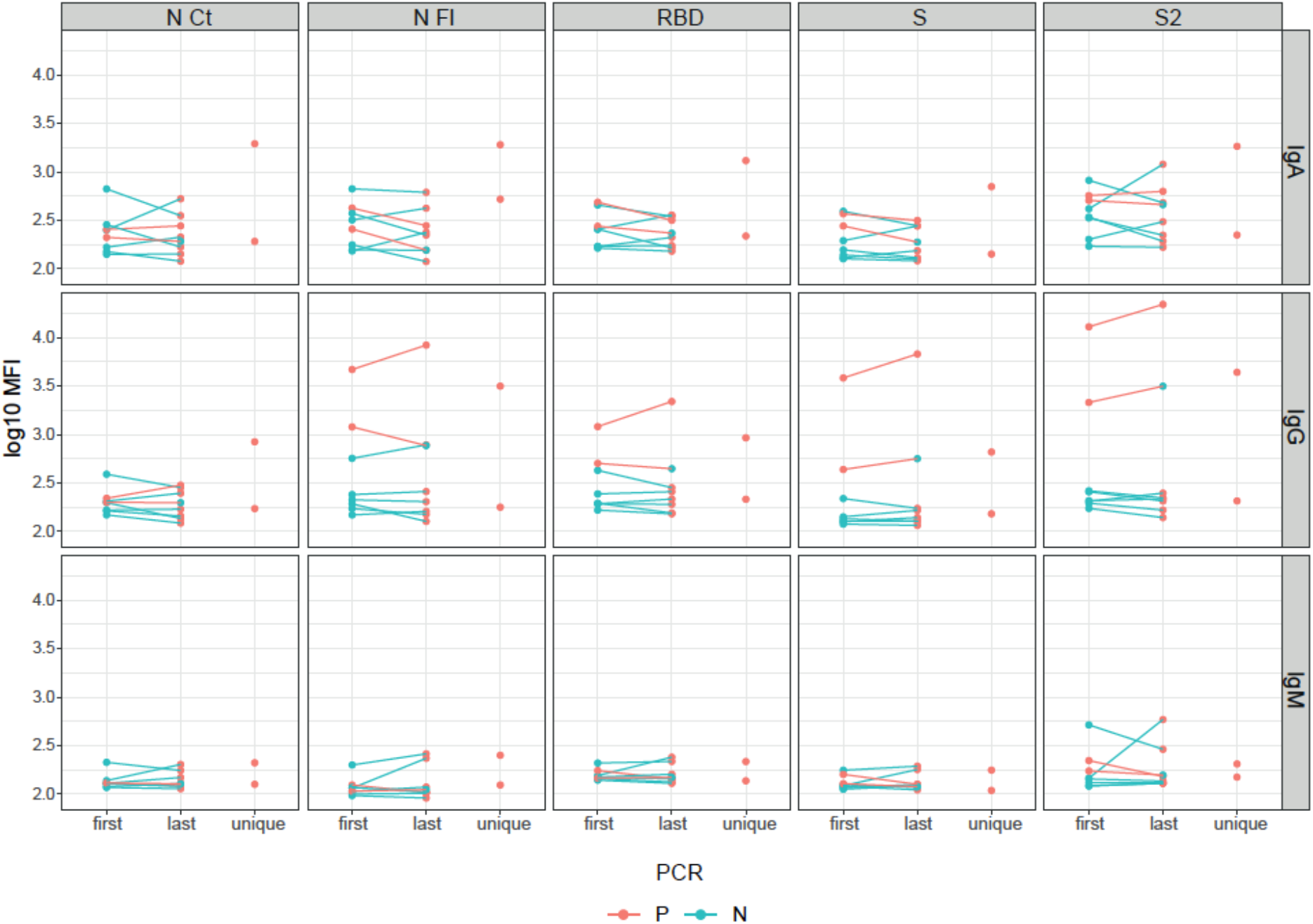
Antibody levels from first to last visits and in unique samples in RT-PCR positives (in orange). RT-PCR positive individuals had higher geometric mean of IgA and IgG levels for most antigens than RT-PCR negative individuals (in green), but not statistically significant (data not shown).

**Figure S3.**
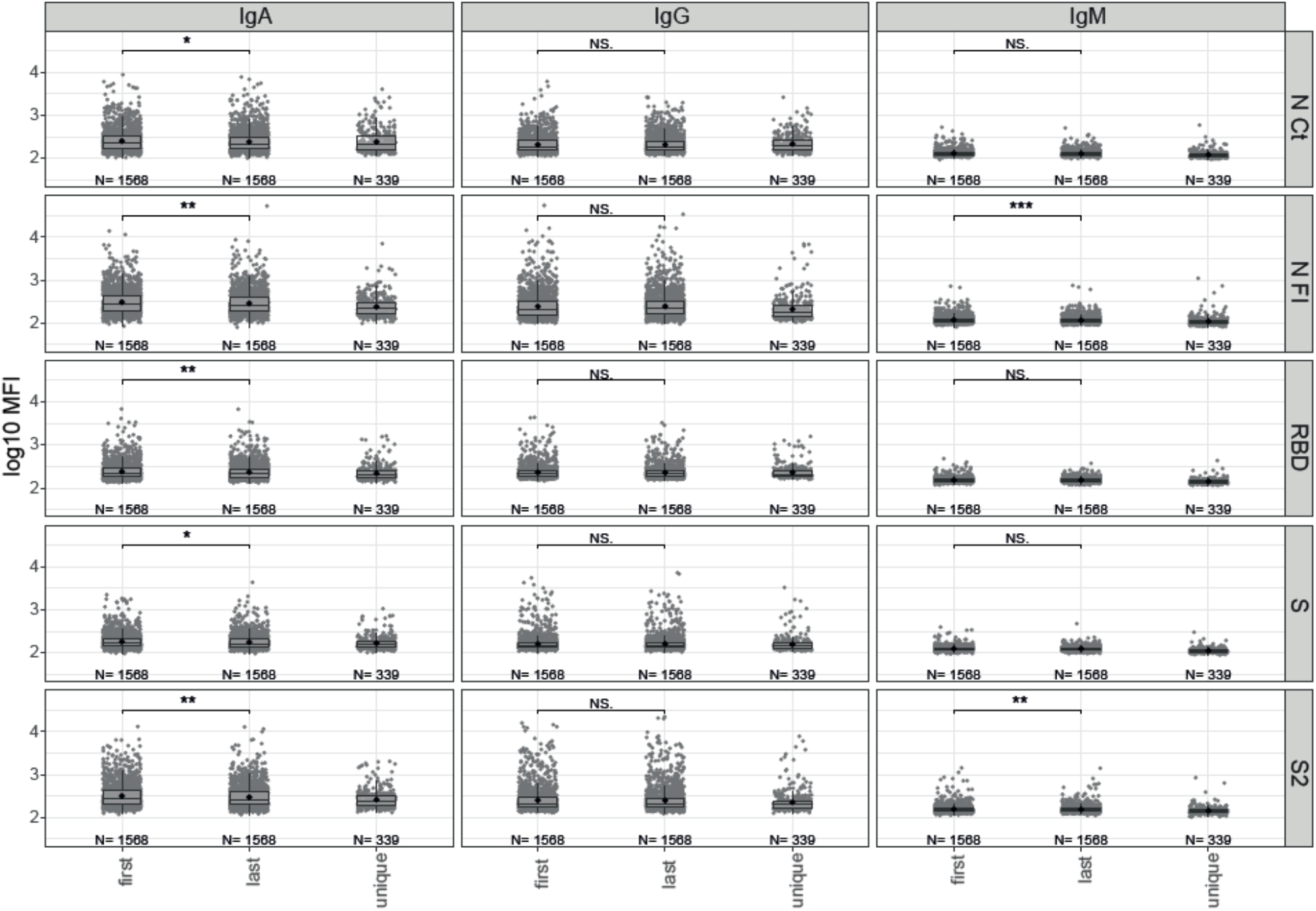
Levels of antibodies at the first, last and single visits. Groups were compared through Mann-Whitney U test. * p ≤ 0.05, ** p ≤ 0.01, *** p ≤ 0.001. NS = not significant.

**Figure S4.**
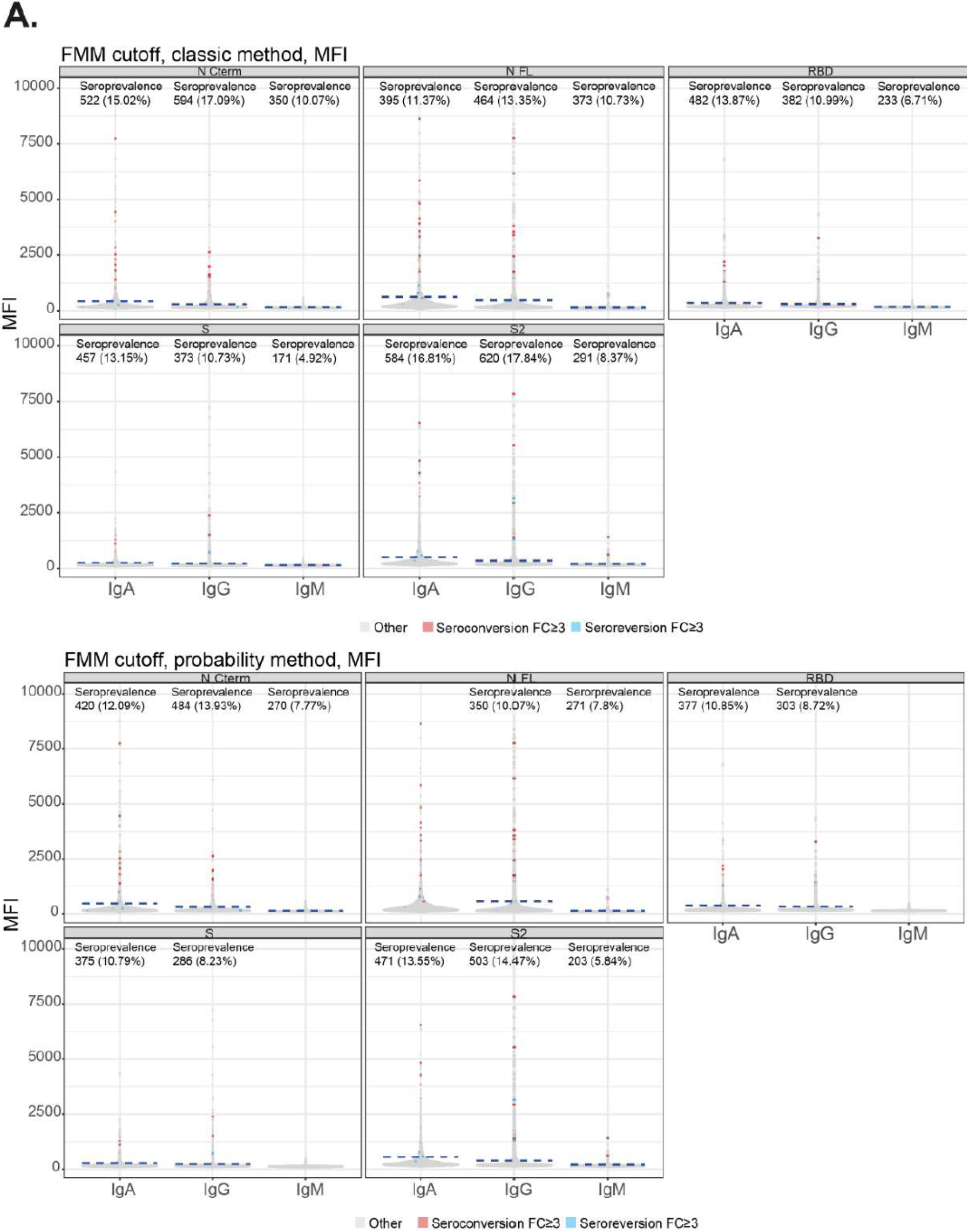
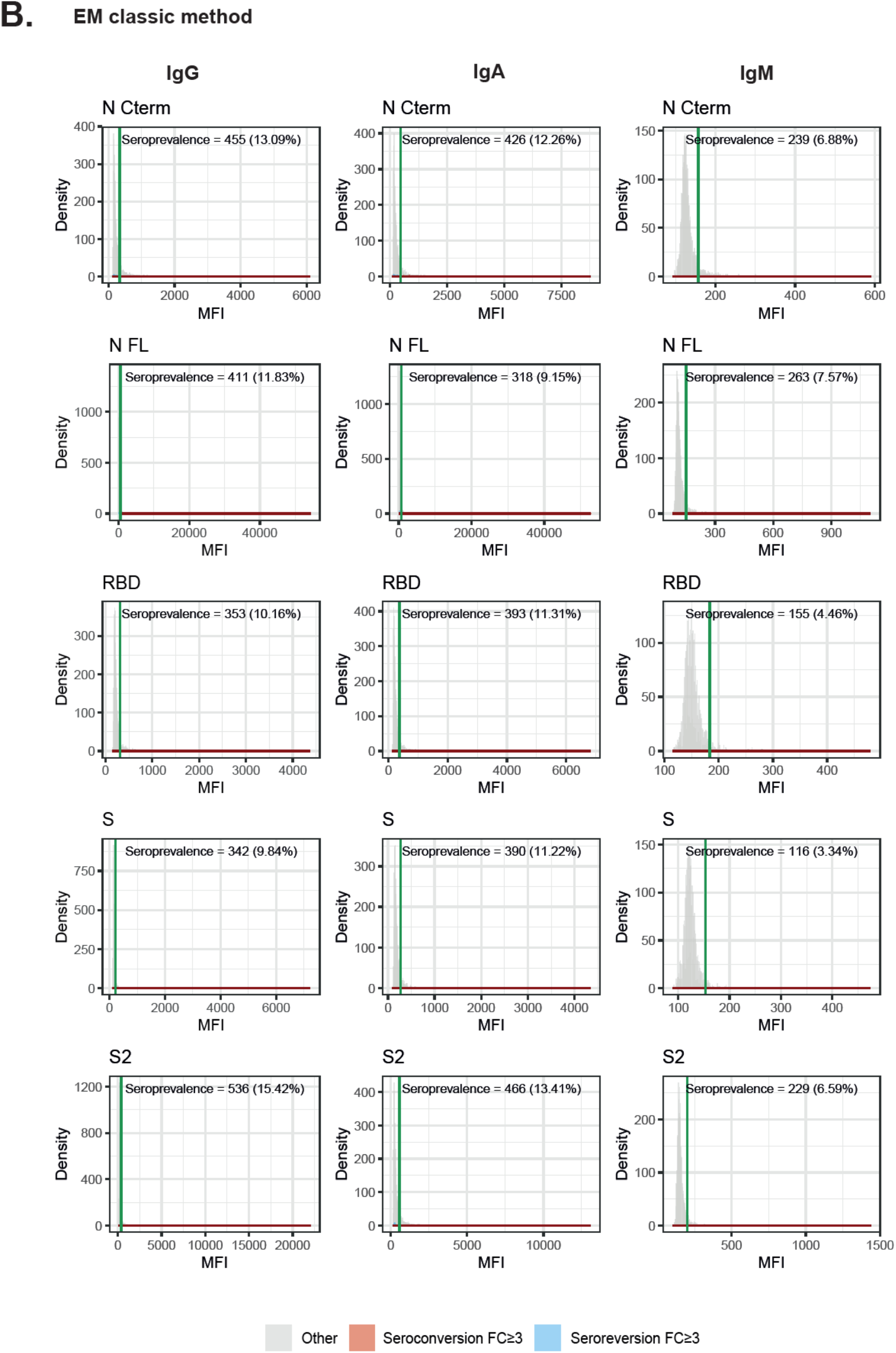
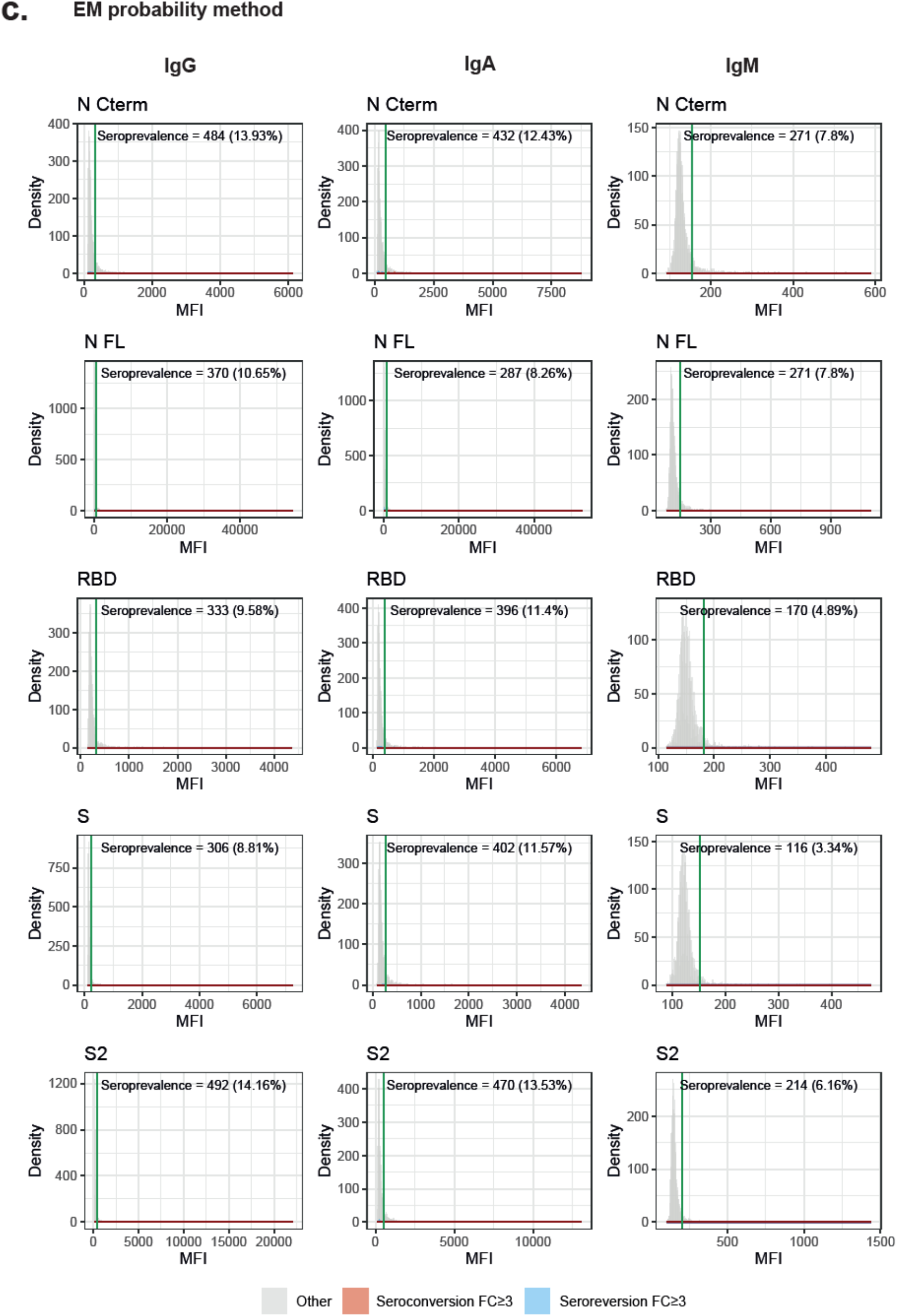
Exploring positivity thresholds and prevalence of anti-SARS-CoV-2 antibody response in pandemic saliva samples by Finite Mixture Models (FMM) and Expectation-Maximization (EM) algorithms in natural MFI scale. In red, samples with ≥3-fold change increase, and in blue samples with ≥3-fold change decrease, in antibody levels in the second visit. SE = seropositivity.

**Figure S5.**
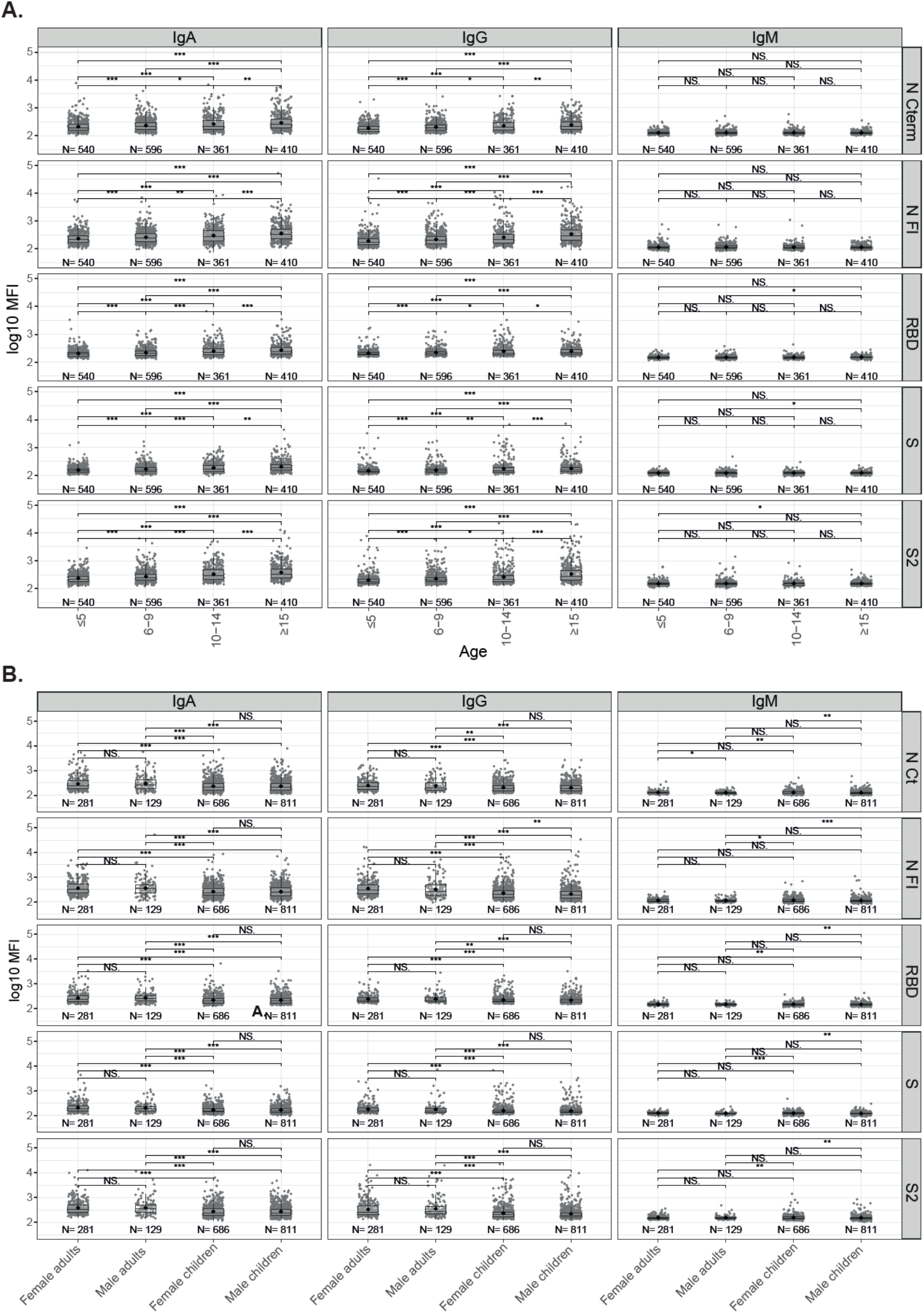
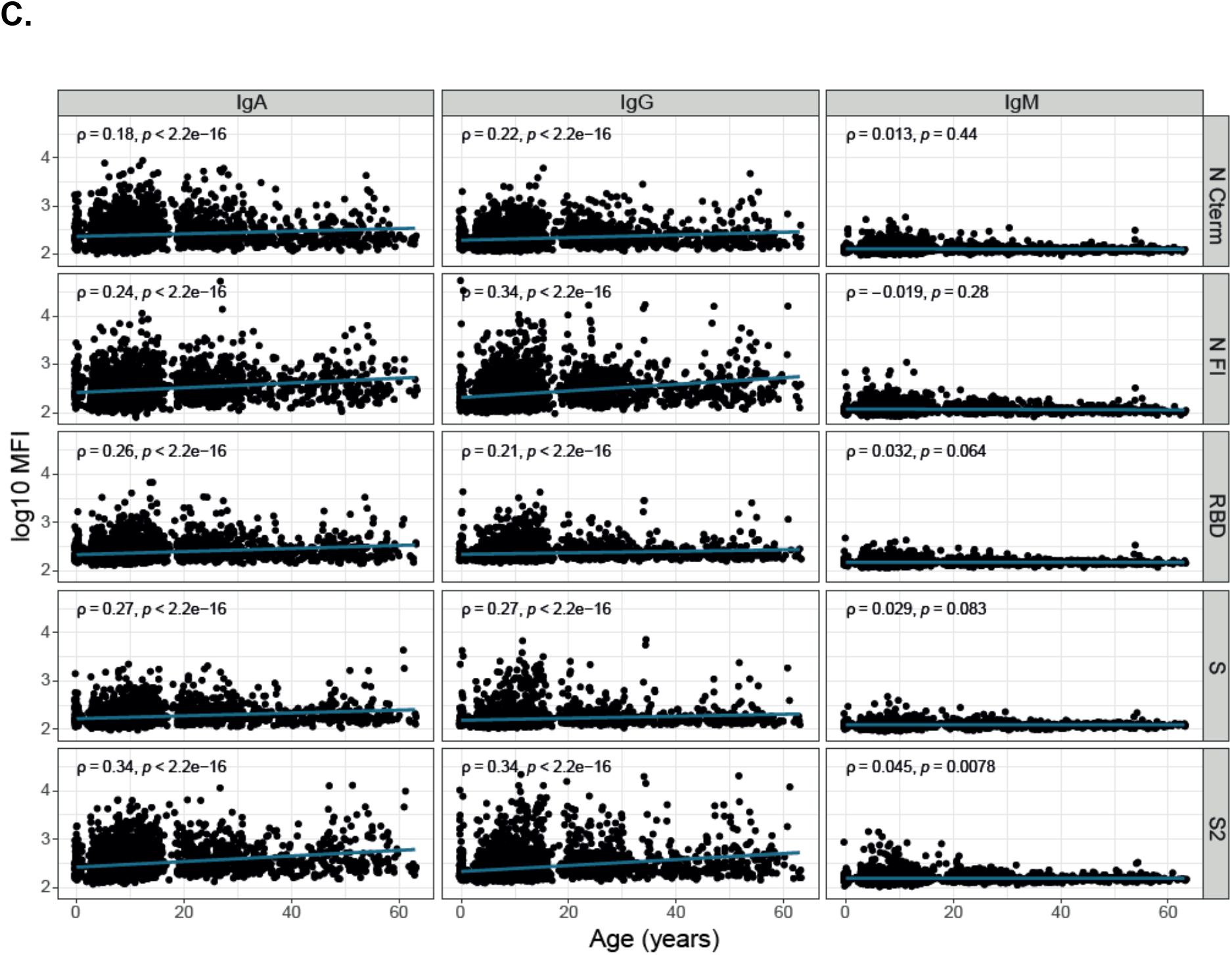
Antibody levels by age. Antibody levels by age category groups in the last and single visits **(A)** and by age and sex **(B)**, compared by Mann-Whitney statistical test, and correlations of antibody levels with age **(C)**. * p ≤ 0.05, ** p ≤ 0.01, *** p ≤ 0.001, NS = not significant.

**Figure S6.**
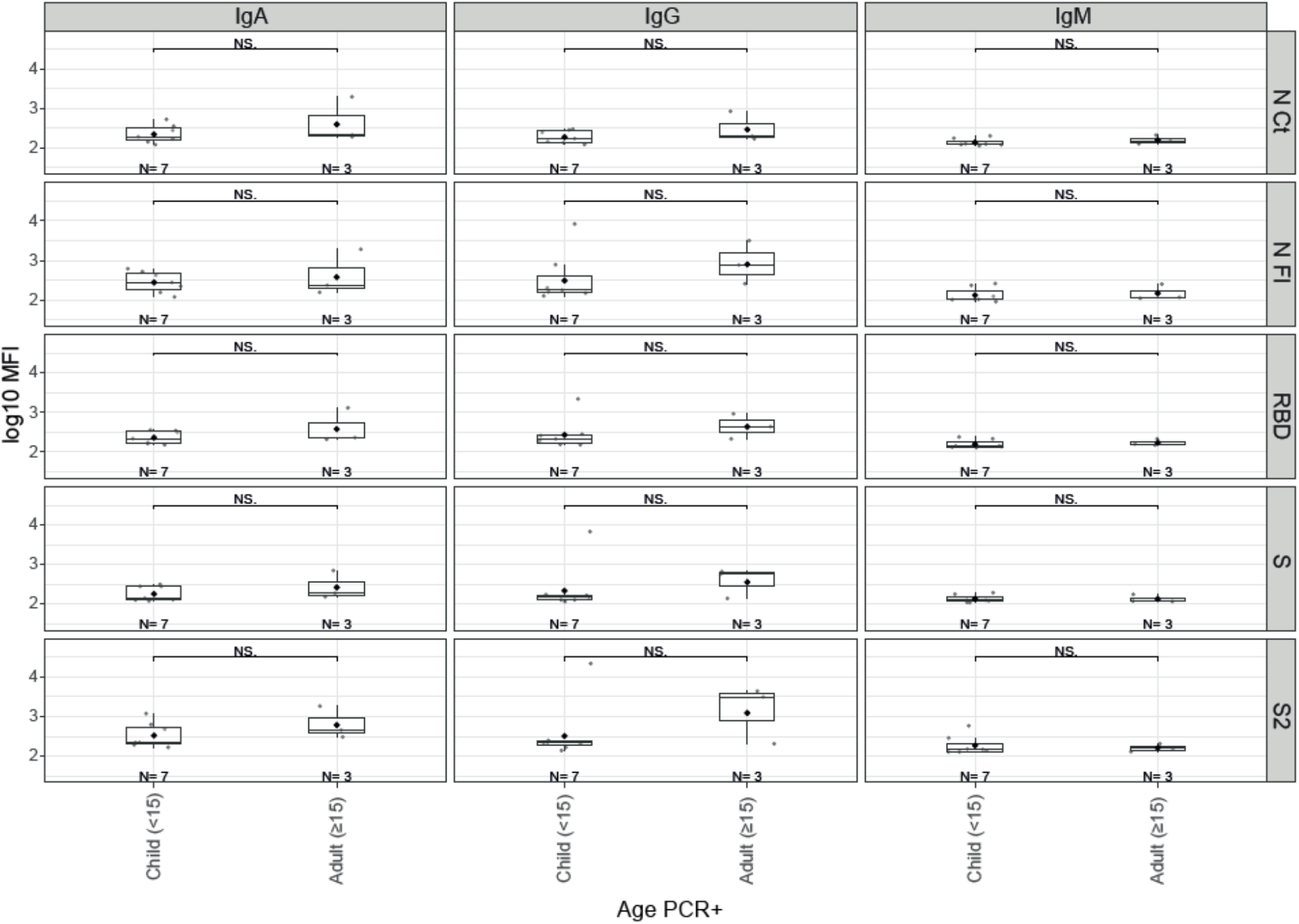
Antibody levels by age in RT-PCR positive individuals. IgA and IgG levels tended to be higher in adults compared to children (<15 years). Groups were compared by Mann-Whitney U test. NS = not significant.

**Figure S7.**
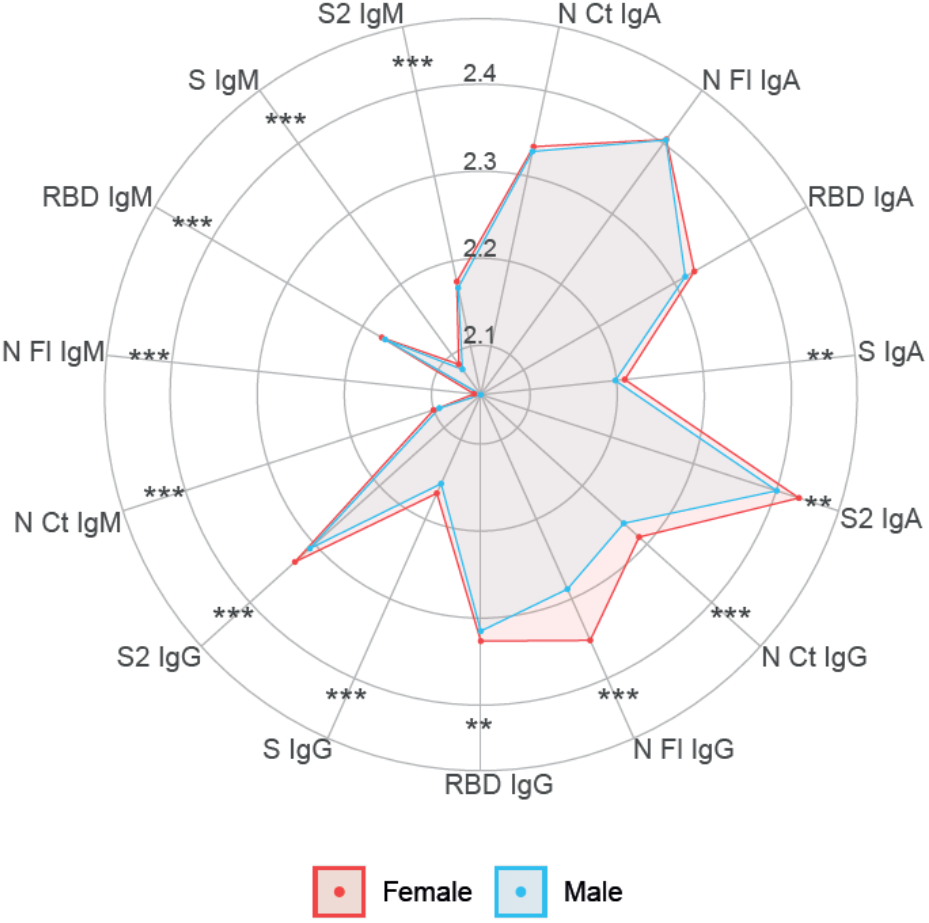
Radar charts of antibody levels by sex. Median antibody levels comparing males (n=940, blue) versus females (n=967, red). Medians were compared through Mann-Whitney U test. * p ≤ 0.05, ** p ≤ 0.01.

